# Partitioning index and non-structural carbohydrate dynamics among contrasting cassava genotypes under early terminal water stress

**DOI:** 10.1101/479535

**Authors:** Luis O. Duque, Tim L. Setter

## Abstract

Cassava (*Manihot esculenta* Crantz.) is a storage root crop of importance in tropical regions where periodic dry season and drought affect performance. Cassava genotypes that differ in performance in ecosystems with various water regimes were subjected to water stress during storage-root initiation and early development. Plants were grown in 50 kg pots in a screen house environment under well-watered and water stress for over a 120-day period. Water stress had a significant effect on most traits analyzed. However, relative water content, partitioning index and non-structural carbohydrates were unaffected. Tolerant genotypes had a higher partitioning index than susceptible genotypes during water stress, associated with a larger number of storage roots initiated and larger storage root biomass, while they were shorter and had less fibrous root biomass. Tolerant lines were indistinguishable from susceptible lines in above ground biomass. These findings suggest that early evaluation of storage root number, partitioning index, and associated traits at an early stage of cassava storage-root development could be an effective approach by which cassava genotypes are screened for favorable drought tolerance response.

## INTRODUCTION

Cassava (*Manihot esculenta* Crantz) is a storage root crop considered a staple food for the rural areas of the tropics because of its inherent adaptation to marginal environments making it an ideal food security and subsistence crop. In the majority of the tropics, it is sown and harvested by smallholder farmers on marginal soils without artificial amendments or controlled irrigation (Cock et al., 1985). It is grown mainly for its starchy tuberous roots and is a key staple food for countless farmers in the tropics (Best and Henry, 1994). Cassava grows reasonably well in low fertility soils and under water deprivation, making it an important staple crop on poverty-stricken marginal lands. Nevertheless, though cassava can endure several months of water stress during its seasonal developmental cycle, water stress still reduces its net biomass production greatly below its maximum yield potential (Connor and Cock, 1981, Connor et al., 1981, El-Sharkawy, 1993, Calatayud et al., 2000, Alves, 2002, Calatayud et al., 2002). The decrease in storage root yield depends on the duration and timing of water deficit and is determined by the sensitivity of a particular growth stage to water stress. Studies have indicated that stress can be particularly damaging the phase from 1 to 5 months after planting, which encompasses stages of storage root initiation and early development. Water deficit during at least 2 months of this period can reduce storage root yield from 32 to 60% (Connor and Cock, 1981, Connor et al., 1981, Porto et al., 1989).

Crops with high and consistent yields and with tolerance to biotic and abiotic stresses are needed. In this regard, selecting cassava for morphological, structural, biochemical, and physiological traits that enhance yield and stress resistance has the potential for raising agricultural productivity (Richards, 2000, El-Sharkawy, 2005).

It is known that drought episodes through current climate variability are a major environmental factor that can limit productivity of crops worldwide. Soil water deficit can be prolonged and chronic in regions with low water availability or random and unpredictable due to changes in weather conditions during the period of plant growth. Thus, understanding crop responses to drought are of great significance and also a fundamental part of abiotic stress breeding schemes and sustainable agriculture (Reddy et al., 2004).

Abiotic stress resistance breeding has fostered many strategies for adaptation to climate change such as, matching phenology to moisture availability using photoperiod-temperature response, increased use of genotypes with known escape and/or avoidance mechanisms during predictable stress events at critical growth and reproductive crop cycles, improved water use efficiency and a reemphasis on population breeding in the form of evolutionary participatory plant breeding to offer a buffer against increasing climate unpredictability (Ceccarelli et al., 2010). Accordingly, improving drought resistance and/or identifying novel drought resistant genotypes in previously known stress resistant crops is an imperative in all plant breeding programs where climate unpredictability is ever present.

In a recent study, Jarvis et al. (2012) predict that cassava will be positively impacted by climate change due to its relative tolerance of high temperatures and water-limited conditions, which are predicted for the African continent. Yet, even though cassava is considered a drought resistant crop, ample diversity exists within the germplasm such that selecting for drought resistance will be beneficial especially under climate change.

Crops, being sessile, have developed specific acclimation and adaptation mechanisms in response to water scarcity. Thorough analysis of these mechanisms will contribute to our knowledge of tolerance and resistance to water stress and assist breeders in the screening and development of novel genotypes resistant to stress.

Overall, crops respond and adapt to drought stress by the induction of various mechanisms such as drought escape or drought resistant mechanisms, with resistance further classified into drought avoidance (i.e. maintenance of tissue water potential) and drought tolerance (Levitt, 1980, Price et al., 2002). Thus, numerous morpho-physiological traits under stress could be used as a proxy or indirectly for selecting for yield under water stress, which in turn can provide drought resistance categorizations in screening studies.

Another category of plant response to drought that is important in crops is altered partitioning and utilization of carbohydrates, which can be limited during stress because of decreased photosynthesis. In cassava, a potential contributor to drought resistant involves its ability to accumulate substantial carbohydrate reserves in its stem, which are slowly remobilized during stress episodes (Duque and Setter, 2013). Another factor is the extent to which a genotype is able to maintain growth of the storage root during stress. In several studies it appears that genotypes well adapted to water stress initiate storage root growth early during development and maintain limited partitioning of carbohydrates toward the storage root during stress (Okogbenin and Fregene, 2002, Okogbenin et al., 2003, Okogbenin and Fregene, 2003).

Considering the large genetic diversity of cassava and its wide distribution within the tropics, the objectives of this study were to: 1) measure and correlate differences between several morpho-physiological traits under early terminal water stress and control conditions; 2) identify and evaluate specific associations between morpho-physiological traits with differences in drought resistance; 3) determine phenotypic variability and genotype relationships and 4) identify traits at an early stage useful for assessing favorable drought resistance response.

## MATERIALS AND METHODS

### Plant Material

The experiment included 15 genotypes that represent a range of landraces and improved cultivars from the International Center for Tropical Agriculture (CIAT) cassava core germplasm collection (Table 1). They were chosen to include lines grown in a diverse range of agro-ecological areas throughout Colombia and Brazil.

**Table 1.** List of the 15 cassava genotypes with their origin, biological and selection status used in this study. CM and GC codes identify types derived from CIAT's cassava breeding program. The remaining genotypes are from the germplasm bank collection.

| Genotype | Code | Synonym | Common name | Origen | Biological status | Selection | Citation* |
| --- | --- | --- | --- | --- | --- | --- | --- |
| BRA 1133 | A | BGM 0491 | Veada 1 | Brazil | Landrace | EMBRAPA/CIAT | (Fukuda et al., 2000; Fukuda et al., 1997) |
| BRA 1243 | B | BGM 0598 | Sapa, Sapa R-16 | Brazil | Landrace | CIAT | (Iglesias et al., 1995) |
| BRA 1435 | C | BGM 0443 | Raimunda | Brazil | Landrace | CIAT | Hernán Ceballos (*pers. comm.*) |
| BRA 255 | D | BGM 0080 | Engana Ladrao | Brazil | Landrace | EMBRAPA/CIAT | (Iglesias et al., 1995) |
| BRA 293 | E | BGM 0549 | Amansa Burro | Brazil | Landrace | EMBRAPA/CIAT | (Fukuda et al., 2000; Fukuda et al., 1997) |
| CG 1141-1 | F | NA | ICA-Costeña | Colombia | Improved line | CIAT | (De Tafur et al., 1997) |
| CM 2772-3 | G | NA | N/A | Colombia | Improved line | CIAT | (CIAT, 1992) |
| CM 3306-4 | H | NA | ICA-Negrita | Colombia | Improved line | CIAT | (De Tafur et al., 1997) |
| CM 4919-1 | I | NA | Corpoica<br>Veronica | Colombia | Improved line | CIAT | (CIAT, 1992) |
| CM 507-37 | J | NA | N/A | Colombia | Improved line | CIAT | (El-Sharkawy et al., 1990) |
| COL 1468 | K | CMC 40 | Mantiqueira | Brazil | Landrace | EMBRAPA/CIAT | (CIAT, 2004; De Tafur et al., 1997) |
| COL 1684 | L | CMC 163 | Charay | Colombia | Landrace | EMBRAPA/CIAT | (CIAT, 2004; De Tafur et al., 1997) |
| PER 183 | M | NA | Eeat 1 | Peru | Landrace | CIAT | (Jaramillo et al., 2005) |
| TAI 8 | N | NA | CMR 246343<br>(Ryg.60) | Thailand | Improved line | EMBRAPA/CIAT | (Cach et al., 2006) |
| VEN 25 | O | UCV 2076 | Querepa<br>Amarga | Venezuela | Landrace | CIAT | (El-Sharkawy et al., 1990) |

### Screen house environment

The experiment was conducted in a screen house at CIAT headquarters in Palmira, Colombia, South America. Approximately 50 stem cuttings (25-30 cm long) of each genotype were disinfected by thermo-therapy (stakes were immersed in 49 **°**C water for 49 minutes) and subsequently immersed in a solution of *Trichoderma spp* (bio-fungicide; 1 kg DW per 55 gallons of water for 10 minutes) and sown individually (one plant per bag) in plastic bags (60 cm in diameter × 40 cm in height; 0.075 m^3^) containing 50 kg sterilized mineral soil : coarse sand (2:1) (15 genotypes × 50 plants = 750 total plants). The soil used was a mollisol (*Aquic Hapludoll*) obtained from the CIAT Palmira station and had the following mean properties at the beginning of the study: pH: 7.65; organic matter: 9.8 g/kg; P (*Brayll*): 3.58 mmol/kg; K: 5.9 mmol/kg; Ca: 65.5 mmol/kg; Mg: 25.1 mmol/kg; Na: 1.5 mmol/kg; cation exchange capacity: 71 mmol/kg; S: 1.14 mmol/kg; B: 0.085 mmol/kg; Fe: 0.35 mmol/kg; Mn: 0.56 mmol/kg; Cu: 0.029 mmol/kg and Zn: 0.72 mmol/kg. Next, plants were placed inside a screen house with corrugated transparent polycarbonate plastic roof (90% light transmission), and side-walls of anti-insect screen, polyethylene monofilament, 266 × 818 µm mesh hole opening size, used to avoid the entrance of whiteflies (*Trialeurodes Aleurotrachelus socialis*) and received manual irrigation to freely drained capacity. At 60 days after planting (DAP), 50 plants of each genotype were randomly assigned to 10 discrete plots. Randomly selected plants had an average height of 160±10 cm and 45±10 unfolded expanded leaves (mean ± SD) at Day 0. One plot of each genotype was then assigned to each of ten blocks and blocks were randomly assigned to either the well watered (WW) or water stressed (WS) treatments (five blocks to each treatment), as described below. Each block contained a complete set of plots representing all genotypes and a given watering treatment, and each plot contained five plants to permit five dates of sampling as described in the next section. Blocks were arranged evenly within the screen house such that all plants within the same block experienced the same screen house lighting, temperature, ventilation, and other environmental conditions. Within each plot, plants were evenly spaced in a grid layout, 0.8 m × 0.8 m, measured from the center of each stem. The distance between plots in all directions was approximately 1.5 m. At this stage (60 DAP), referred to as Day 0, two water treatments were imposed: (i) control (plants were irrigated every other day until field capacity) and (ii) water stress (irrigation was withheld and soil was allowed to dry for the duration of the experiment. All plants were maintained inside the screen house for the duration of the experiment.

### Growth parameters

Plants within a plot were randomly assigned to five sampling dates to assess development stage effects in response to WW and WS treatments. The first sampling date was at 60 DAP (referred as to Day 0), at which point irrigation in the WS treatment was stopped. The second sampling date was 15 days after Day 0 (referred as to Day 15) and in successive manner, Day 30, Day 45, and finalizing at Day 60. At each sampling date, a set of morpho-physiological traits were recorded (Table 2 and described in detail in later sections).

**Table 2.** Morpho-physiological traits recorded in cassava genotypes subjected to both well watered and water stress conditions

| **Trait designation** | **Code** | **Unit** | **Description of Trait** |
| --- | --- | --- | --- |
| Plant height | PH | cm | Height of the main stem from soil level to the uppermost fully expanded leaf |
| Leaf relative water content | RWC | % | Leaf water content at sampling to that present at full turgor |
| Storage root | SR | g FW | Fresh storage root weight of individual plants |
| Above ground biomass | AGB | g FW | Fresh biomass (stems, petioles and leaves) weight of individual plants |
| Fibrous root | FR | g FW | Fresh fibrous root weight of individual plants |
| Number of storage roots | #SR | - | Total number of storage roots of individual plants |
| Partitioning index | PI | - | Ratio of storage root weight to total biomass |
| Leaf retention | LR | % | Measured from length of the first intact leaf-petiole to the uppermost apical meristem and plant height |
| Leaf abscisic acid | L-ABA | pmol g<sup>-1</sup> DW | Leaf abscisic acid concentration measured from fully expanded mature leaves |
| Leaf glucose | L-GLC | μmol g<sup>-1</sup> DW | Leaf glucose concentration measured from fully expanded mature leaves |
| Leaf total sugars | L-TS | μmol g<sup>-1</sup> DW | Sum of leaf glucose and sucrose concentrations measured from fully expanded mature leaves |
| Leaf starch | L-STR | μmol g<sup>-1</sup> DW | Leaf starch concentration measured from fully expanded mature leaves |
| Stem abscisic acid | S-ABA | pmol g<sup>-1</sup> DW | Stem abscisic acid concentration measured from middle portion of stem |
| Stem glucose | S-GLC | μmol g<sup>-1</sup> DW | Stem glucose concentration measured from middle portion of stem |
| Stem total sugars | S-TS | μmol g<sup>-1</sup> DW | Sum of stem glucose and sucrose concentrations measured from middle portion of stem |
| Stem starch | S-STR | μmol g<sup>-1</sup> DW | Stem starch concentration measured from middle portion of stem |
| Root abscisic acid | R-ABA | pmol g<sup>-1</sup> DW | Root abscisic acid concentration measured from mature root tips |
| Root glucose | R-GLC | μmol g<sup>-1</sup> DW | Root glucose concentration measured from mature root tips |
| Root total sugars | R-TS | μmol g<sup>-1</sup> DW | Sum of root glucose and sucrose concentrations measured from mature root tips |
| Root starch | R-STR | μmol g<sup>-1</sup> DW | Root starch concentration measured from mature root tips |
| Soil water content | SM5 | θ | Volumetric soil water content measured at 5 cm depth |
| Soil water content | SM25 | θ | Volumetric soil water content measured at 25 cm depth |

### Plant height and leaf retention

Plant height (PH) was measured as the distance from the soil surface at the base of the main stem to the uppermost fully expanded leaf. In cassava, leaf abscission advances in a highly predictable pattern starting at the lowest stem node and advancing upward, with retained leaves in the apical section of the stem and branches. To ensure equal comparison among genotypes differing in plant height, leaf retention was calculated by measuring the length from the first intact leaf-petiole to the uppermost apical meristem on the main stem (height of the stem containing retained leaves, HRL) and PH according to the following expression:

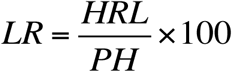

### Soil water content

Volumetric soil water content (*θ*, m^3^×m^-3^) was measured in the first 0-5 cm (SM5) and 20-25 cm (SM25) soil layers on each plant using a ThetaProbe Soil Moisture Sensor (model ML2x; Delta-T Devices, UK). A set of three-pronged waveguide rods made of stainless steel, 20 cm long and 3.0 mm in diameter, was inserted horizontally in each soil layer and allowed to equilibrate. A total of two measurements per soil layer were taken and averaged.

### Yield components

At each sampling date, plant biomass and its components were measured including aboveground biomass fresh and dry weight (AGB), storage root fresh and dry weight (SR), fibrous root fresh and dry weight (FR), number of storage roots (#SR) and fresh weight partitioning index (PI). A plant from each WW and WS plot was harvested at Day 0, 15, 30, 45, and 60. Storage roots were defined as roots >5 mm diameter. Partitioning index of storage roots (PI) was measured as the ratio between storage root fresh weight and total biomass expressed in the following equation:

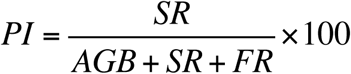

### Non-structural carbohydrates and abscisic acid

At each sampling date (Day 0, 15, 30, 45, and 60) a total of three tissue samples were collected from each plant. Three leaf disks (diam. = 0.635 cm) were sampled from three mature fully expanded leaves and another three from the three uppermost folded immature (expanding) leaves to form a composite sample.

Afterwards, cylindrical stem plug samples from the middle third of the shoot system were obtained utilizing a 3 mm diameter cork borer. Three fibrous root tips were sampled by cutting about 1 cm of small healthy portions from the new root growth to form a composite sample per plant. All plant tissues were sampled at Day 0, 15, 30, 45, and 60 between 1100 and 1400 hours. Sampled tissue was immediately immersed in 300 µL of ice-chilled (0°C) 80% methanol and then stored at −20°C until further use. All leaf measurements were expressed on an area basis; stem and root measurements were expressed on a residual dry weight basis. Sucrose, glucose and, starch were measured on all plant tissue extracts using the peroxidase/glucose oxidase (PGO) method, with invertase and amyloglucosidase to hydrolyze sucrose and starch to sugars, as described by (Ober et al., 1991) and modified by (Setter et al., 2001).

Prior to hormone analysis, leaf tissue was first separated into fractions based on hydrophobicity using reverse phase C_18_ chromatography, as described by Setter and Parra (2010). Briefly, the method involves separation with C_18_ mini-columns (solid phase extraction columns; model DSC-18 SPE-96, Supleco, Bellefonte, PA) in a 96-well vacuum apparatus. Samples were loaded in 30% (v/v) methanol with 1% v/v acetic acid, and ABA was eluted with 65% methanol with 1% acetic acid. Abscisic acid (ABA) levels were determined using an enzyme-linked immunosorbant assay (ELISA) as described by Setter et al. (2001).

### Relative Water Content (RWC)

RWC expresses the quantity of water in a tissue relative to the absolute quantity of water which the plant would need to achieve complete saturation and is used to assess leaf water status in water stress scenarios (Gonzalez and Gonzalez-Vilar, 2003). Measurements of RWC were performed between 1100 and 1400 hours on each of the sampling dates. A composite sample of 3 leaf discs (diam. = 1.9 cm) was sampled from three mature fully expanded leaves. Leaf RWC was determined with the following equation (Barrs and Weatherley, 1962, Smart and Bingham, 1974):

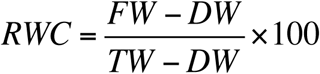

Leaf fresh weight (FW) was determined immediately after sampling, whereas turgid weight (TW) was determined by soaking the composite leaf samples in distilled water in test tubes for up to 12 hours at 20°C. After soaking, leaf samples were quickly and carefully blotted dry with Kimwipe (Kimberly-Clark, Roswell, GA USA) tissue paper in preparation for determining turgid weight. Dry weight (DW) was assessed after oven drying leaf samples at 60°C for 48 hours.

### Statistical Analysis

All data were subjected to analysis of variance (ANOVA), correlation, principal components analysis (PCA) and cluster analyses, which was carried out using R (version 2.15, R Foundation for Statistical Computing, http://www.r-project.org/). The ANOVA model contained the following factors: genotype (G), watering treatment (T), G x T interaction, sampling date (S), S x T, S x G, and residual. All sources of variation were considered fixed. PCA, derived biplot and cluster analysis were conducted for genotypes, environments and drought resistance indices, using squared Euclidean distance as the proximity measure and incremental sum of squares as the grouping strategy (Ward, 1963). This was performed to interpret relationships among selection criteria, compare genotypes on the basis of morpho-physiological differences and to identify genotypes or groups of genotypes with a measureable level of drought resistance.

## Results

### Analysis of Variance (ANOVA)

Water stress significantly (p<0.05) affected 17 out of 22 traits at the last sampling on Day 60 (Table 3). Water stress decreased growth of plant height, aboveground biomass (AGB), storage root dry mass (SR), number of storage roots (#SR), fibrous roots (FR), and plant height (PH). As expected, soil moisture was less in water stress, and ABA in leaves, stems, and roots were elevated; however, leaf RWC in water stress did not differ from controls. However, partitioning index (PI), which reflects the proportion of partitioning into storage roots, differed between genotypes, but was not altered by water stress. Genotype effects were significant for a large number of traits, 14 out of 22 (*P*<0.01 and *P*<0.001) indicating a high level of genetic variability. These included morpho-physiological traits such as leaf retention (LR), leaf starch (L-STR), leaf abscisic acid (L-ABA), stem glucose (S-GLC), stem total sugars (S-TS), stem starch (S-STR), stem abscisic acid (S-ABA) and root total sugars (R-TS) (Table 3). Significant genotype × treatment interactions were found for volumetric soil water content at 25 cm depth (SM25), L-ABA, S-GLC, S-TS, S-ABA, and R-TS indicating variable performance of genotypes in both growing conditions (Table 3).

**Table 3.**
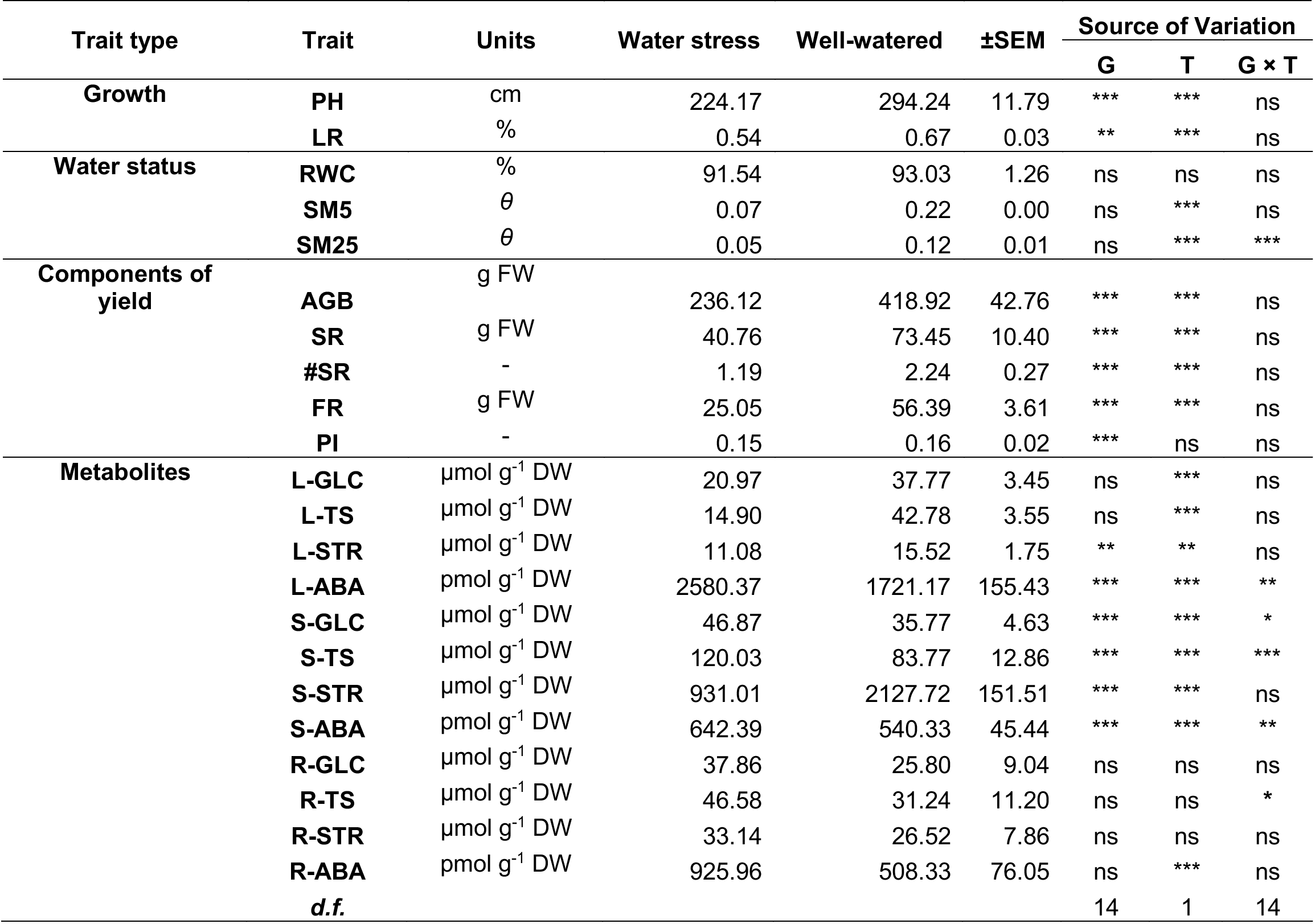
Summary of analysis of variance (ANOVA) for genotype (G), watering treatment (T) and genotype × treatments interaction (G × T) for traits analyzed for sampling Day 60. Shown are the means, pooled SEM, ***d.f***.:degrees of freedom, and significance of effects: *** *p* < 0.001; ** *p* < 0.01; * *p* < 0.05; ns: non-significant

### Phenotypic correlations

To further understand the interrelationship of measured traits under WS conditions, a phenotypic correlation matrix was constructed using data from Day 60 and presented in Table 4. SR yield was significant and correlated with AGB (*r* = 0.68**), RWC (*r* = 0.6*), LR (*r* = 0.80***), PI (*r* = 0.76***), #SR (*r* = 0.67**), L-STR (*r* = 0.58*), S-GLC (*r* = 0.54*), and S-STR (*r* = 0.66**). However, under WW conditions, SR yield was correlated with #SR (*r* = 0.92***), PI (*r* = 0.77**), and LR (*r* = 0.52*). In addition, SR yield under control conditions presented non-significant positive correlations with AGB, PH, and S-STR. In regards to other important morpho-physiological trait associations, PI was highly correlated #SR (*r* = 0.61*), and SSTR (*r* = 0.57*) under WS conditions.

**Table 4.**
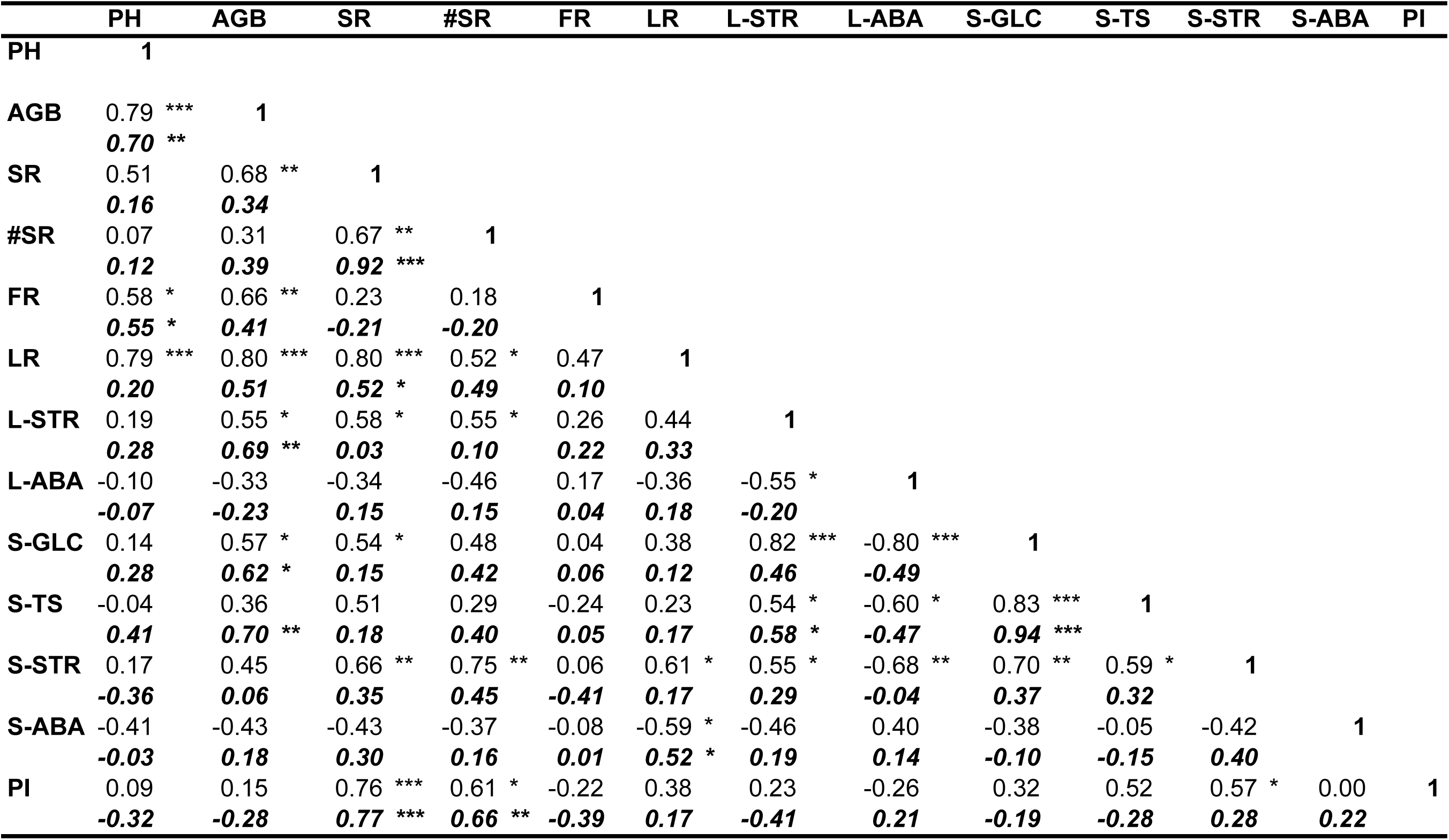
Phenotypic correlation coefficients between traits for 15 cassava genotypes under water stress and well watered conditions taken at Day 60. Values in bold and italics represent well watered conditions. *** *p* < 0.001; ** *p* < 0.01; * *p* < 0.05

Correspondingly, under WW conditions, PI was significant and correlated with SR, #SR, and R-STR, and non-significant with, LR, and S-STR. LR was significant and correlated with PH, AGB, #SR, L-GLC, L-TS, S-STR and S-ABA under WS, while under WW conditions LR was non-significant with leaf, stem or root non-structural carbohydrates traits. In addition, S-GLC was both highly correlated with L-STR (*r* = 0.82***) and negatively correlated with L-ABA (*r* = −0.80***) under WS conditions.

### Principal Components and two-way hierarchical cluster analysis

Principal components analysis (PCA) for data under water stress at Day 60 was used to provide a reduced dimension model that would indicate measured trait differences among the 15 cassava genotypes under water stress. Traits that showed a significant genotype effect were used as input variables for this analysis.

The first three principal components explained 79% of the observed variation under stress (Table 5). Specifically, PC1 explained 48% of the variation and showed the largest loading correlation values with SR, LR, S-GLC and S-STR. PC2 explained 20% of the variation and showed high loading correlation values with FR, PH, and AGB. Lastly, PC3 explained 10% with high loading correlation values including PI and S-GLC.

**Table 5.** Loading scores and variance for PC1, PC2 and PC3 of the principal component analysis (PCA) for traits under water stress and well-watered conditions for 15 cassava genotypes

| **Traits** | **Water stressed** |  |  | **Well-watered** |  |  |
| --- | --- | --- | --- | --- | --- | --- |
|  | **PC1** | **PC2** | **PC3** | **PC1** | **PC2** | **PC3** |
| **PH** | 0.198 | 0.471 | 0.037 | 0.256 | -0.236 | 0.358 |
| **AGB** | 0.307 | 0.322 | -0.137 | 0.440 | -0.142 | 0.182 |
| **SR** | 0.348 | 0.047 | 0.342 | 0.254 | 0.419 | 0.119 |
| **#SR** | 0.287 | -0.112 | 0.279 | 0.310 | 0.372 | 0.000 |
| **FR** | 0.108 | 0.489 | -0.064 | 0.088 | -0.296 | 0.437 |
| **LR** | 0.324 | 0.287 | 0.155 | 0.271 | 0.197 | 0.342 |
| **L-STR** | 0.307 | -0.064 | -0.299 | 0.334 | -0.167 | -0.008 |
| **L-ABA** | -0.269 | 0.253 | 0.328 | -0.135 | 0.210 | 0.393 |
| **S-GLC** | 0.324 | -0.215 | -0.352 | 0.386 | -0.117 | -0.327 |
| **S-TS** | 0.252 | -0.348 | -0.112 | 0.414 | -0.151 | -0.278 |
| **S-STR** | 0.333 | -0.168 | 0.090 | 0.187 | 0.275 | -0.345 |
| **S-ABA** | -0.226 | -0.140 | 0.219 | 0.113 | 0.226 | 0.244 |
| **PI** | 0.221 | -0.235 | 0.606 | -0.023 | 0.499 | -0.005 |
| **Eigenvalue** | 6.297 | 2.561 | 1.344 | 4.1511 | 3.2759 | 1.9393 |
| **% of variation** | 48.44 | 19.7 | 10.33 | 31.932 | 25.199 | 14.918 |
| **Cumulative %** | 48.44 | 68.14 | 78.48 | 31.932 | 57.131 | 72.048 |

To further classify the 15 genotypes under WS based on the PCA, a biplot of PC1 and PC2 and a two-way hierarchical cluster using all 13 PC scores were constructed (Figs. 1 and 2). PC1 shows the majority of traits with positive loading values except for L-ABA and S-ABA, which scored negative. PC2 shows positive loading values for FR, PH, AGB, LR and SR (yield and components of yield traits) and negative loading values for #SR, L-STR, S-STR, S-GLC, PI and S-TS (non-structural carbohydrate traits).

**Figure 1.**
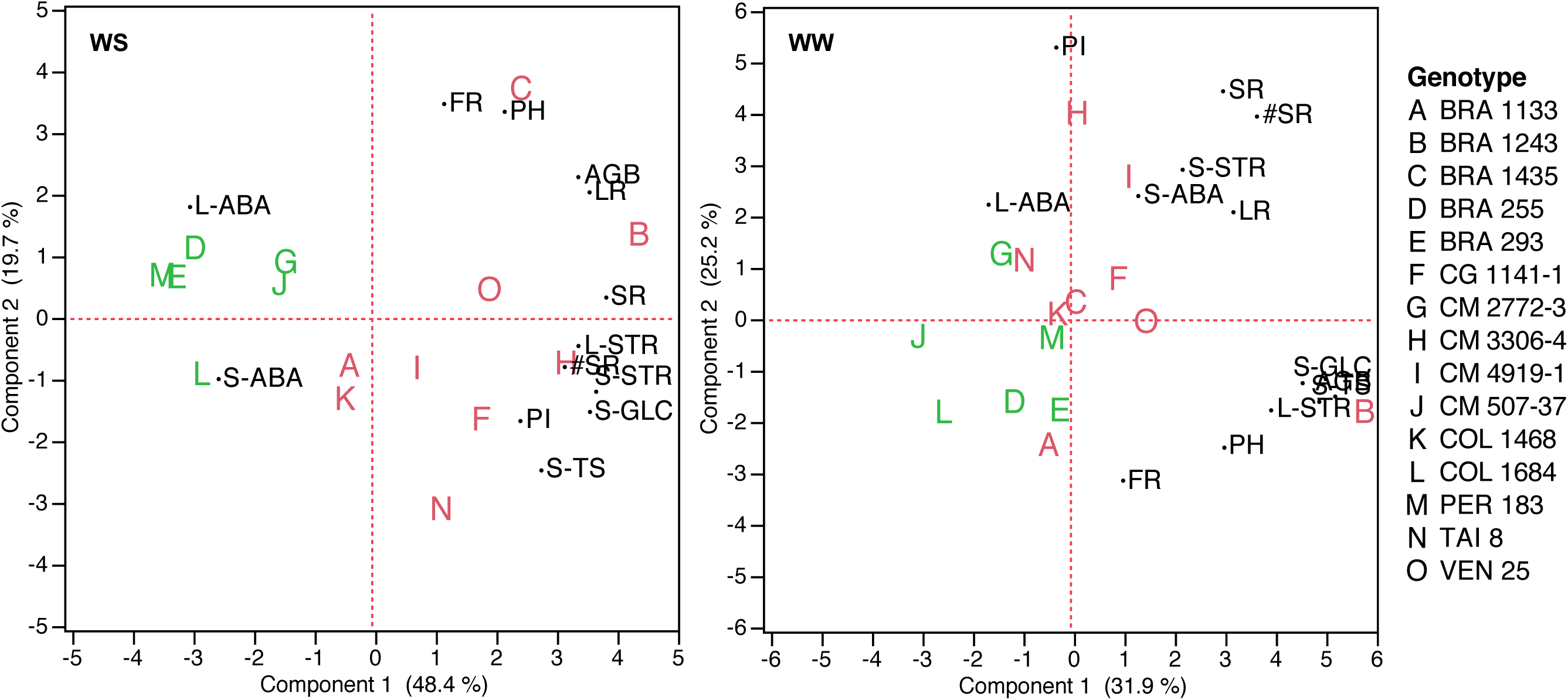
Biplot of principal components analysis for traits under water stress (WS) and well watered (WW) conditions of 15 cassava genotypes. Color-coded genotypes show clustering by trait grouping.

**Figure 2.**
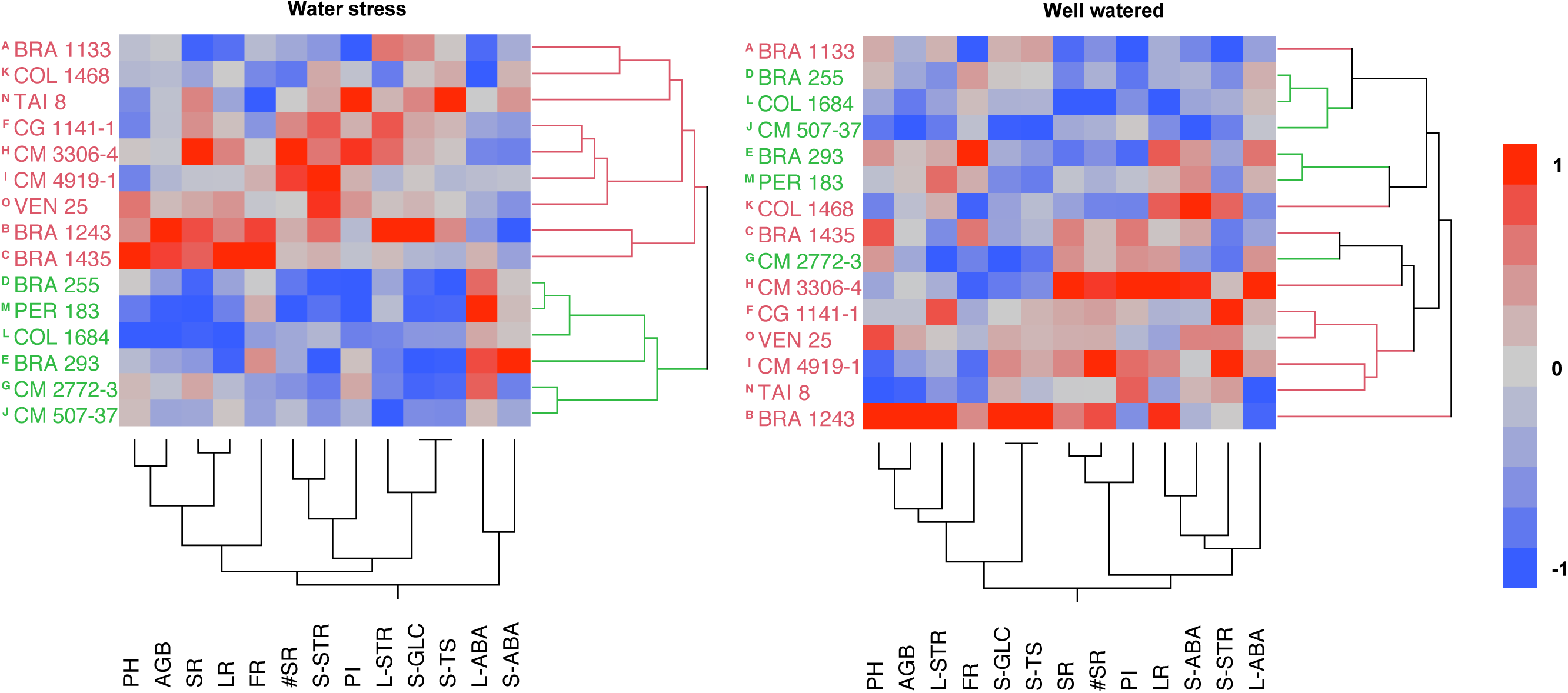
Two-way hierarchical cluster dendrogram based on genotypes and traits for water stress (WS) and well-watered (WW) conditions. Color-coded genotypes show clustering by trait grouping. Heat map shows the values of all the data colored across its range value.

Genotypes B, C, F, H, I, N, and O were identified as ones with moderate to high storage yield (PC1) and moderate to high components of yield (PC2) under WS. While genotypes A, K, D, E, G, J, L, and M presented low SR yield (low PC1) and high L-ABA and S-ABA (high PC2).

The two-way cluster analysis generally confirmed the result of the PCA. For example, under WS, genotypes were classified into two separate clusters (Fig. 2). The first group included genotypes (A, B, C, F, H, I, K, N, and O; in red), which had moderate to high PC1 and PC2. The second group clustered genotypes (D, E, G, J, L, and M; in green), which presented moderate PC1 and PC2 scores. Interestingly, genotypes A and K were grouped in cluster 1 possibly on the basis of non-structural carbohydrates and not on SR yield, PI of LR.

### Regression analysis relationships and temporal dynamics

The relationship between several morpho-physiological traits and SR yield were plotted to observe behavioral properties both under WS and WW conditions (Fig. 3). The linear regression of all genotypes’ SR yields under WW and WS conditions are shown in Fig. 3A. These results indicate that genotypes with high SR yield under control conditions also placed high in when subjected to water stress (R^2^ = 0.75). This possibly indicates that a high SR yield potential under optimal conditions could result in high SR yield potential under stress for selected elite genotypes. Figures 3B, C, D, E and F shows the relationship between LR, L-ABA, S-STR, PI, and L-STR with SR yield for all genotypes both WW and WS conditions. These results indicate a positive trend for LR (R^2^ = 0.64), S-STR (R^2^ = 0.44), PI (R^2^ = 0.59) and L-STR (R^2^ = 0.33) and a negative trend for L-ABA (R^2^ = 0.11) for genotypes under water stress. Specifically, these results showed that genotypes with higher SR yield under stress displayed higher LR, S-STR, PI and L-STR and lower L-ABA. Similarly, selecting genotypes with high values for these traits could result in high yield potential under WS conditions.

**Figure 3.**
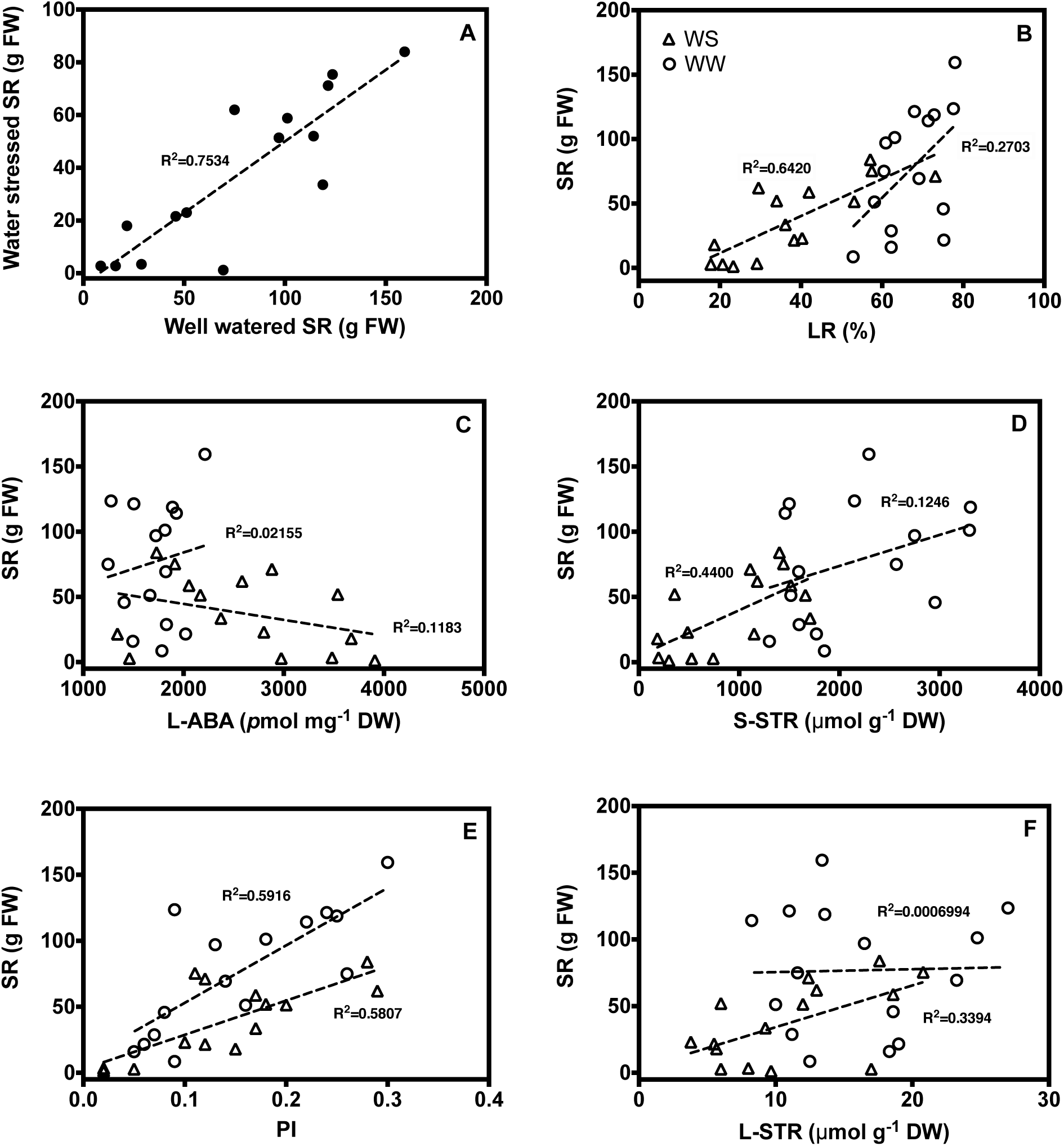
Regression relationship between storage root yield under stress and control conditions (A); leaf retention (B); leaf abscisic acid; stem starch (D); partitioning index (E); and leaf starch (F) for each genotype under well watered and stress conditions. Open circles are genotypes under well watered conditions (WW); open triangles are genotypes under water stress conditions (WS).

To determine the time frame over which the water deficit exerted effects, water status and morphological growth measurements were made at 15-day intervals. For this analysis, cluster 1 (C1; genotypes A, B, C, F, H, I, K, N, and O) and cluster 2 (C2; genotypes D, E, G, J, L, and M) were used to represent temporal dynamics of morpho-physiological traits (Fig. 4), and non-structural carbohydrates and abscisic acid (Fig. 5).

**Figure 4.**
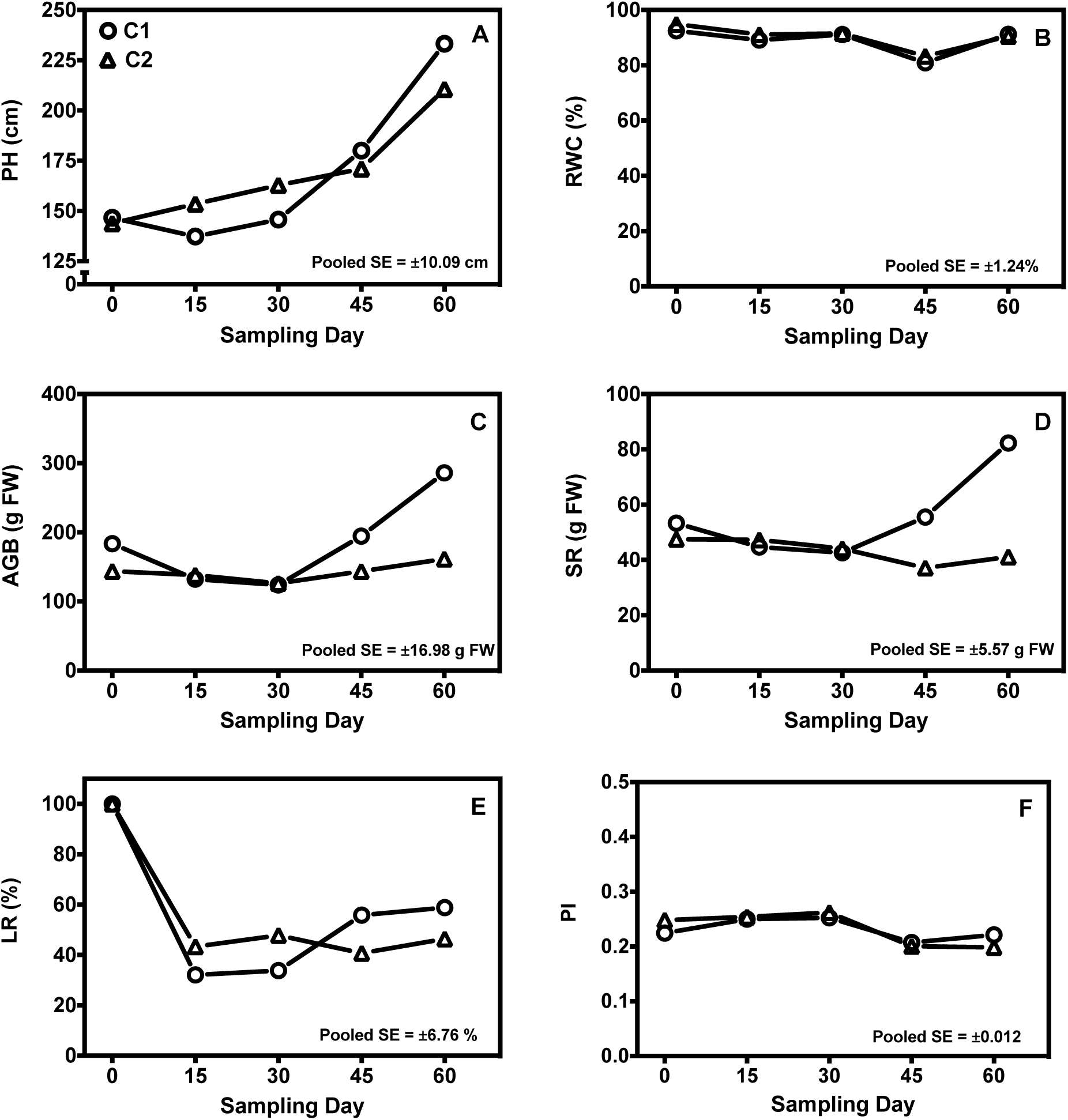
Temporal dynamics of morpho-physiological traits of clustered cassava genotypes under water stress over a period of 60 days. ±Pooled SE: pooled standard error; n = 75 plants per sampling day. Plant height (A); relative water content (B); above ground biomass (C); storage root (D); leaf retention (E); and partitioning index (F)

**Figure 5.**
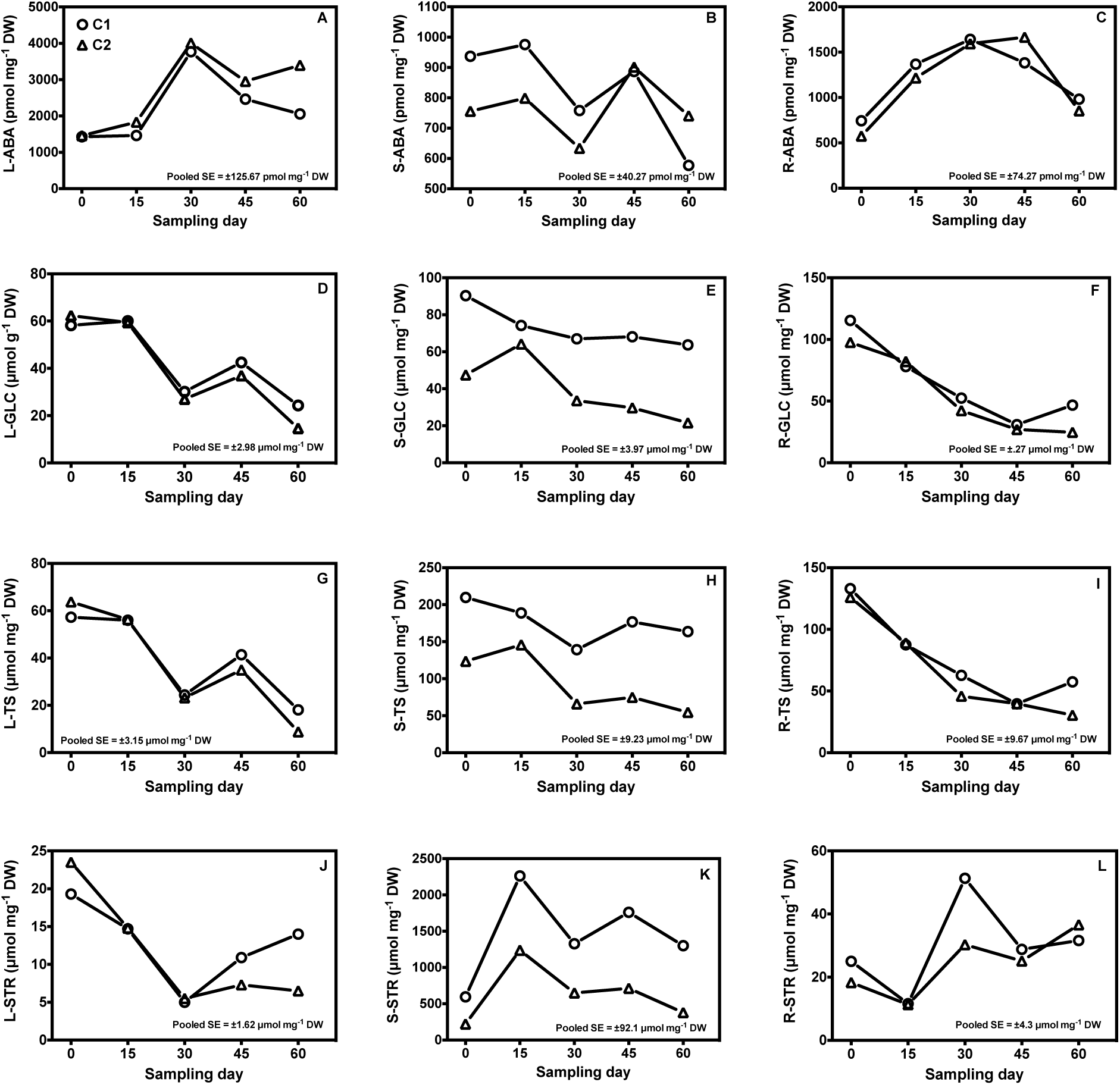
Temporal dynamics of abscisic acid (ABA) and non-structural carbohydrates (NSC) in mature leaves, stems and fibrous roots of clustered cassava genotypes under water stress over 60 days. L-ABA (A); R-ABA (C); S-ABA (B), L-GLC (D); S-GLC (E); R-GLC (F); L-TS (G); S-TS (H); R-TS (I); L-STR (J); S-STR (K); and R-STR (L). ±Pooled SE: pooled standard error; n = 75 plants per treatment/sampling day. Cluster 1 is composed of genotypes: A, B, C, F, H, I, K, N, and O. Cluster 2 is composed of genotypes: D, E, G, J, L, and M.

Under stress, both C1 and C2 presented a gradual increment in plant height (PH) with genotypes in C1 favoring a higher PH by Day 60 (Fig. 4A). Despite depletion of soil water, relative water content (RWC) of upper-canopy leaves remained indistinguishable from well-watered controls throughout the experiment (Fig. 4B). Nevertheless, growth data indicated that the water stress treatments were exerting effects on plants as early as Day 15.

Aboveground biomass fresh weight (AGB), and total root fresh weight (SR+FR) were similar both in C1 and C2 at about Day 30. Afterwards, genotypes in C1 showed an positive increase both in AGB and SR+FR until Day 60, while genotypes in C2 remained low (Figs. 4C and 4D). In addition to organ growth, another early event in stress response was leaf abscission.

Water stress stimulated substantial leaf abscission so that by Day 15 leaf retention (LR) fell to ~30% in C1 and ~40% in C2 (Fig. 4E). Both clusters subjected to water stress had a minimum percent of LR by Day 30. Interestingly, percent LR in cluster 1 increased until the end of the experiment. This was due to shoot growth (Figs. 4C, 4D) and associated production of new leaves at the apical meristem. The measured value for LR was a function of countervailing leaf abscission and new leaf formation. Partitioning index (PI), showed an initially increase in both C1 and C2 but severely decreased by Day 45 with a slight increase in C1 by Day 60 (Fig. 4F).

Figure 5 presents the results of both non-structural carbohydrates (NSC) and abscisic acid (ABA) in leaves, stems and root segments. Under water stress, abscisic acid (ABA) concentration in leaves was significantly higher in C2 than in C1 by Day 45 and remained high by Day 60 (Fig. 5A). In stems, ABA concentration in C1 was higher than C2 and tended to decline from Day 0 to Day 45. However, by Day 60, C2 showed higher SABA when compared to C1. This phenomenon was associated with sampling young green stems at early stages, and more woody and starch-filled specimens at later stages (Fig. 5B). In fibrous roots, water deficit increased ABA levels in both clusters by Day 15, and remained high throughout the remainder of the experiment (Fig. 5C). These data indicate that the timing of water stress sustained growth inhibition and ABA accumulation for the period from Day 15 to 60. In regards to non-structural carbohydrates, sugar and starch concentration in leaves decreased substantially in response to WS in both clusters (Figures 5D, 5G and 5J). Glucose was the most abundant non-structural carbohydrate in leaves and by Day 60 both L-GLC and LTS were slightly higher in C1 when compared to C2. Leaf starch (L-STR), decreased in both clusters by Day 30, however by Day 60, L-STR increased in C1 while C2 remained low (Fig. 5J). A similar difference was observed in stem NSC (Figs. 5E, 5H, and 5K). Overall, stem glucose (S-GLC), total sugars (S-TS) and starch (S-STR) in cluster 1 were significantly higher when compared to cluster 2. Starch was the predominant storage carbohydrate in stems, so this difference represents considerably more storage carbohydrate in genotypes grouped in cluster 1 than genotypes in cluster 2. In fibrous roots, root glucose (R-GLC) and total sugars (R-TS) decreased under water stress starting by Day 15 and were indistinguishable between both clusters. However, by Day 60, C1 presented an increase in both root NSC when compared to C2 (Figs. 5F and 5I). Root starch (R-STR) presented an overall increased tendency in both clusters by the end of the experiment (Fig. 5L).

## Discussion

Cassava responds to water deficit in the form of changes in various physiological and morphological processes (Duque and Setter, 2013). These morpho-physiological changes have been considered as important adaptation mechanisms for cassava to resist drought (Okogbenin et al., 2013). However, the physiological basis of genetic variation in water stress response and its association with yield and related traits is not clear in cassava. In this study, a small but diverse panel of cassava genotypes was evaluated for differences in drought resistance, interrelationships between traits and storage root yield and temporal dynamics of stress response.

The water stress treatment was begun at a relatively early stage of cassava development, 60 days after planting the propagation stakes (Day 0), and imposed for a further 60 days, which coincides with the timing of storage root initiation and early bulking (Deoliveira et al., 1982, Okogbenin and Fregene, 2002). Some studies have indicated that cassava storage root development is especially sensitive to drought and photosynthesis-limiting shade stress at this stage, such that storage root yield is severely affected by a relatively short-term stress, whereas stress at later stages of root bulking is less damaging to yield (Deoliveira et al., 1982, Aresta and Fukai, 1984). Other studies have indicated that cassava’s development is set back by early drought stress, but is capable of recovering (Baker et al., 1989, El-Sharkawy and Cadavid, 2002).

Overall, water stress had a significant effect on most traits analyzed (Table 3). However, relative water content (RWC), partitioning index (PI) and root non-structural carbohydrates were not affected. Leaf RWC under stress remained at values similar to controls throughout the experiment in the full set of 15 diverse genotypes (Fig. 4B). Maintenance of RWC occurred while soil water content was depleted and leaf growth was inhibited. This behavior is thought to be caused by decreased transpiration due to leaf abscission and acute sensitivity of stomata to minor decreases in leaf water potential (*Ψ*_W_) during periods of water stress or low humidity and high transpiration demand (El-Sharkawy and Cock, 1984, Itani et al., 1999). The current study supported cassava's characterization as a *drought avoider*, in the sense that it downwardly adjusts its water loss to avoid exposing its tissues to extremely low water status.

As discussed above, the target water stress episode was imposed at 2 MAP for 60 days without re-watering (terminal stress) in order to study the effects of water deficit during the time of active root bulking and growth. In cassava, partitioning index (PI) is the ratio of storage root yield as a fraction of the total plant biomass measured at 4–5 months in contrast to harvest index (HI) which is typically measured at 12 MAP. In our study, PI under stress presented a proportional reduction both in storage root yield and aboveground biomass when compared to well watered controls but statistically PI was not affected by water stress and had similar values to the well-watered controls (Table 3; Fig. 3E). In addition, the correlation between SR yield and PI under stress was significant (Table 4). Overall, our results showed that genotypes with higher PI under well-watered conditions also placed high under stress and suggests that an important component of greater water stress resistant in cassava is a genotype’s tendency for storage root initiation and sustained PI and storage-root development during stress.

Under field conditions with prolonged water stress, some studies have observed that while cassava produces less total biomass, it increases its partitioning index into storage roots (i.e., harvest index) (Connor et al., 1981, El-Sharkawy et al., 1992, El-Sharkawy and Cadavid, 2002). This has been explained as a consequence of water stress inhibition of stem biomass accumulation, leaf abscission, and vigorous leaf regrowth during periods of renewed rainfall. In addition, other studies have shown a positive correlation between PI (at 7 MAP) and HI (at 12 MAP), PI stability across many environments and potential genetic gain with the use of PI (i.e. HI) in cassava breeding programs (Mutegi-Murori, 2009, Olasanmi, 2010, Okogbenin et al., 2013). These findings support the breeding approach of selecting for early-bulking genotypes, particularly for drought-prone environments (Deoliveira et al., 1982, Okogbenin and Fregene, 2002, Okogbenin et al., 2013).

Leaf retention (LR) or conversely leaf abscission is a trait that has been extensively studied in cassava (Ike and Thurtell, 1981, Ramanujam, 1990, Lenis et al., 2006, Duque and Setter, 2013). In the current study, LR was substantially decreased by WS, contributing to the overall decrease in accumulation of aboveground biomass (Table 3; Fig. 3B and Fig. 4E). While cassava genotypes grouped in cluster 1 (C1) and cluster 2 (C2) was not discernibly different in LR temporal dynamics (Fig. 4E), overall, there does exist genotypic differences between individuals (Fig. 3B). Specifically, genotypes that scored high LR under stress and control conditions also scored high for SR yield under both conditions. This is consistent with a study by Turyagyenda et al. (2013) of a drought resistant and a sensitive line, and by (Lenis et al., 2006) involving 1350 cassava clones subjected to a 5-month dry period towards the end of the growth cycle, where yield was correlated with leaf retention. In addition, studies indicate that percent leaf retention in cassava is the net result of leaf abscission and new leaf formation (Connor and Cock, 1981, Duque and Setter, 2013).

The principal component analysis (PCA) is one of the most effective techniques that have been used in different area of sciences (Johnson and Wichern, 2007). The aim of PCA is to reduce the potentially large dimensionality of data space (observed variables) to smaller intrinsic dimensionality of feature space independent variables. In the present study, under water stress the first principal component (PC1) had higher positive correlations with SR, S-STR, S-GLC and LR, and negative correlations with leaf and stem ABA while the second principal component (PC2) presented positive correlations with FR and PH. (Table 5). These results indicate that under water stress, higher SR yields are achievable with more leaf retention, sustainable stem non-structural carbohydrates and lower organ-specific ABA synthesis. This result is consistent with a study performed by Duque and Setter (2013) where a basal rate of leaf growth continued and stem and storage roots maintained relatively high starch concentrations and contents per organ until final harvest under stress. In addition, stems gradually lost starch and had sufficient reserves to serve as a prolonged source of remobilized carbohydrate during stress. Organ specific ABA synthesis (explained in PC1) and fibrous root growth (FR; explained in PC2) also play an important role in understanding cassava’s drought resistance among genotypes and its relationship to SR yield under stress. Studies performed by Alves and Setter (2000), showed that cassava rapidly accumulated ABA under stress and was completed reversed one day of with re-watering. Also, ABA concentrations were differentially correlated with genotypes suggesting that genetic variability could exist within contrasting cassava genotypes. Within our study, leaf ABA varied greatly among genotypes evaluated both under water stress and control conditions (Fig. 3C). Yet, the best performing genotypes for SR yield under stress (cluster 1) produced less L-ABA than genotypes in cluster 2 (Fig. 5A).

Estimation of cassava’s fibrous root system under contrasting environments is difficult. However, it has been shown that although growth of fibrous lateral roots was impaired by water stress, main root elongation to deeper regions was only modestly decreased (Duque and Setter, 2013). In addition, other studies have shown ample genotypic differences exist in fibrous root weight and length after 2 to 5 (WAPS) under water deficit (CIAT, 1994).

Taking these observations into account, individual selection or clustering of genotypes (notably C1 and C2; Fig. 1) with high PC1 and high PC2 may result in genotypes with superior drought resistance. Based on these two components and according to the distribution of cassava genotypes on the biplot (Fig. 1), genotypes C, B, and H with high PC1 and PC2 values may be suggested as superior genotypes under stress. Furthermore, the widespread distribution of genotypes on the biplot (Fig. 1), which was later confirmed by the cluster analysis (Fig. 2) indicated a large genetic variation in the studied plant population in response to water stress.

Among organ-specific NSC temporal dynamics under stress, L-GLC and L-TS in both clusters progressively decreased until Day 60, however L-STR recovered in C1 genotypes and remained higher when compared to C2 genotypes by the end of the experiment (Day 60) (Figs. 5D, 5G, and 5J). Several studies have shown that during active photosynthesis excess CHO accumulates and stored as starch in leaves as a temporary reserve and are the principal component of dry mass accumulated in mature leaves, whereas sugars are transported to developing sink organs were little or no photosynthesis takes place (Basu and Minhas, 1991, Lorenzen and Ewing, 1992, Saeedipour and Moradi, 2011). Under stress, photosynthesis declines fomenting a synthesis and decrease of leaf starch to sugars and remobilizes sugars to different plant organs to sustain metabolic activities. In cassava, a similar trend has been observed, however a study by Duque and Setter (2013) indicted that remobilized sugars did not contribute to stress acclimation. Instead, they declined coincident with deceases in transpiration rate and were an indicator of the stress effects on carbon balance.

Conversely, L-STR increased after Day 45 in genotypes grouped in cluster 1 when compared to C2. This singularity is probably due to increases in photosynthesis in newly formed and retained apical leaves (LR) in C1 genotypes. Contrary to sugar and starch levels in leaves, overall stem NSC levels were high in cluster 1 when compared to cluster 2 under stress (Figs. 5E, 5H, and 5K). Specifically, stems accumulated large quantities of starch (S-STR; Fig. 5K). In studies by Duque and Setter (2013) cassava plants at an early stage of storage-root growth had levels of starch in stems that represented 35% of the plant reserve carbohydrate and 6% of whole plant biomass. In the current study, S-STR accumulated under stress by Day 15 and slightly decreased until Day 60. This trait contributes to the ability for remobilization of starch from stems to other plant parts during extended periods of drought (Duque and Setter, 2013, Okogbenin et al., 2013).

Studies of several crop species have indicated that utilizing stem reserves under stress can improve carbon balance under photosynthate-limiting conditions (Blum, 2005, Reynolds and Tuberosa, 2008). Specifically, in grain crops, soluble carbohydrate reserves in the stem at the time of anthesis may contribute to superior performance under drought stress (Kumar et al., 2007), and are associated with improved yield potential in field environments where temporary storage helps plants cope with short-term stresses that limit photosynthate supply (Shearman et al., 2005). The long growing season in cassava, and growth in environments that often span long dry seasons, increase the likelihood that reserve carbohydrate storage and remobilization is a valuable trait in this crop.

In order to ensure the efficient and effective use of cassava germplasm, its characterization is essential. In the present study, 13 out of the 22 variables measured under water stress contributed to the apparent variation among the cassava genotypes examined (Table 3). The greater part of the variation was accounted for SR yield, yield components and stem NSC. Although the cluster analysis grouped genotypes with greater morphological similarities under stress, the clusters did not necessarily include germplasm from the same origin or biological status (Table 1). Though our study did not center its attention on the association between morphological characteristics and geographical origin of the cassava genotypes, merit exists in developing ideotype-breeding strategies for specific environmental scenarios. Altogether, progress in adopting the ideotype model for breeding cultivars adapted to various environmental conditions has been demonstrated in cassava (El-Sharkawy, 2006).

The results from our work also indicate that phenotypic assessment of SR yield stability under water stress and control conditions, an important breeding objective, can be effectively determined by PCA of selected morpho-physiological traits. Cassava genotypes grouped in cluster 1 may be combined with breeding lines exhibiting high yield potential under well-watered conditions. However, a thorough genotypic analysis in combination with physiological studies is prerequisite to establish whether these genotypes are comparable (i.e. deploy the same resistance mechanisms). Provided, different mechanisms are identified, there is potential for recombining these for further improvement.

In summary, the current study identified several attributes that sustain storage root *status quo* under water stress and potential genotypic differences between genotypes examined. Cluster 1 and 2 genotypes differed in reserve carbohydrate accumulation: cluster 1 had higher starch levels in leaves and stems during stress, while fibrous roots had higher total sugars towards the end of the experiment. These findings could be due to organ-specific events related to development and carbohydrate remobilization, and deserve further study to relate them to underlying processes and the genes responsible for the effect. Consistent with cassava’s characteristic water stress avoidance, the current studies showed that RWC remains high despite terminal water deficit and leaf retention data indicated that genotypes with less leaf abscission presented higher storage root yield under stress. In addition, organ-specific ABA temporal dynamics showed that genotypes with less leaf ABA accumulation displayed sustained NSC synthesis and storage root yield. This stomatal response was apparently due to genotypes having a high degree of stomatal sensitivity, as leaves of cluster 1 genotypes accumulated a less leaf ABA than genotypes in cluster 2 by the end of the experiment.

Our results indicated that even though there was a penalty in early storage root yield under stress, several genotypes placed high both under water stressed and control conditions. In addition, genotypes that differed significantly for storage root yield and morpho-physiological traits both under stress and control conditions indicate considerable genetic variation within the germplasm evaluated. This suggests that a high storage root potential under optimum conditions could result in improved storage root yield under water stress conditions. Thus, the use of early SR yields, as an indirect selection criterion for a drought-prone environment based on optimum conditions could be efficient.

Following this train of thought and of particular importance for yield improvement in stress environments, the current work showed that several genotypes had a higher storage root partitioning index than others during the stress. This result was associated with genotypes having a consistently larger number of storage roots initiated during stress and larger storage root biomass, while plants were shorter and had less fibrous root biomass. Correspondingly, several genotypes presented higher aboveground biomass from other under stress. Thus, the observed genotypic differences in storage root partitioning index, which were measured at an early stage of development and which were associated with performance under stress, suggest that early evaluation of storage root biomass could be an effective method by which cassava genotypes are screened for favorable drought resistance response.

## Acknowledgements

The authors would like to thank Alfredo Alves and Martin Fregene for their assistance in initiating this work and for advise on cassava germplasm. We thank Grace Liu and Cheryl Chiang for lab assistance. We thank Juan Carlos Pérez and Fernando Calle for field and screen house assistance.

